# Do tropical birds avoid evasive prey? An experimental study in Ecuadorian avian communities

**DOI:** 10.64898/2026.05.29.728647

**Authors:** Erika Páez, Héctor Cadena-Ortiz, Edison Ocaña, Marianne Elias, Violaine Llaurens

## Abstract

The effect of predator behaviours on the evolution of colour pattern has been extensively studied in chemically defended prey but much less so in evasive prey, although similar selection regimes might be at play. Most previous work on the recognition and learning of colour patterns by predators has relied on experiments with few model predator species, preventing a proper assessment of how natural predator communities shape the evolution of prey signals. Here, we investigate predation on evasive iridescent blue *Morpho helenor* butterflies by wild avian communities in a tropical forest, using artificial butterflies exposed to natural bird assemblages attracted to insect-light traps in the field. We presented five prey types: evasive (local and exotic Morpho), cryptic, palatable-control and unpalatable-control. We recorded attacks from 43 different bird species, and compared attack latency, attack order and predation exerted by different birds on different prey types. Most birds avoided evasive and defended prey, but attack responses differed depending on their levels of specialization towards insect prey. Avoidance of conspicuous coloured prey was more prevalent in specialist flight-feeding, invertivorous birds, whereas more opportunistic treehunter and frugivorous guilds had weaker discrimination. Exotic and local evasive iridescent blue Morphos experienced similar predation rates, suggesting a generalization of iridescent blue signals associated with evasiveness by most birds. Finally, contrasting selection on ventral vs. dorsal patterns was detected in *M. helenor*, where cryptic ventral surfaces were exposed to attack by ground-foraging birds, whereas conspicuous iridescent blue coloration displayed during flight is associated with reduced attack rates. By revealing potential differences in the responses of specialist and generalist predators, this study highlights the importance of accounting for diversity of predator traits and behaviours when investigating the evolution of aposematism, mimicry, and other anti-predator adaptations.

## INTRODUCTION

The evolution of morphological and behavioural traits in species submitted to strong predation pressure is jointly shaped by the local communities of both predator and prey species. Local predators can indeed learn from previous experiences and therefore tend to avoid unpalatable prey (Skelhorn and Rowe 2006), as well as evasive prey (Linke et al 2022; Páez et al 2021). The attack decisions depend on the communities of available prey (Lindström et al 2004) and shape the evolution of traits in prey species. Predator behaviours triggered by visual cues may lead to either apostatic selection promoting increased variability in cryptic colour pattern in palatable prey (Bond and Kamil 2002) or positive density-dependent selection limiting colour pattern variation in defended prey (Nokelainen et al 2024). Furthermore, many predators also display neophobic behaviour, thereby avoiding attacks on prey with unfamiliar appearance (e.g., in insectivore birds, (Greenberg and Mettke-Hofmann 2001). Therefore, to identify the selection exerted by visual predators on the evolution of coloration in different prey species requires to characterise (i) the local diversity of predators attacking the prey species and (ii) the predator’s attack decision regarding local defended and palatable prey vs. exotic ones. Lepidoptera have been an excellent model of study of predation pressures since they display a wide variety of aposematic or cryptic wing colour patterns (Ruxton et al 2004). In this study, we used the palatable species *Morpho helenor* to explore how the behaviour of natural predators may shape the evolution of aposematism and mimicry in an evasive species. This species is distributed in Central and South American tropical forests (Blandin and Purser 2013). It displays bright iridescent blue colour on the dorsal side of the wings and brownish coloration on the ventral side, leading to a flashing effect during flapping flight (Murali 2018). This peculiar dorso-ventral contrast, combined with fast erratic flight (Le Roy et al 2021), likely enhances the escape abilities of these butterflies when attacked by bird predators (Vieira-Silva et al 2024). Experiments with wild jacamars show low attack success on these butterflies in flight, whereas resting *M. helenor* individuals with closed wings are eaten, confirming their palatability (Pinheiro and Campos 2019). Furthermore, the survival of *M. helenor* may stem not only from the direct effect of these escape abilities during attacks, but also from indirect effects due to predator learning and avoiding to attack evasive prey (Pinheiro et al 2016). Experiments with naïve birds indeed showed that predators can learn to associate wing colour patterns with difficulty of capture, and therefore stop attacking prey with high evasive capacities (Páez et al 2021). Such learning behaviour may then favour the convergent evolution of similar colour patterns across evasive prey facing the same predator community, similarly to the Müllerian mimicry process (Ruxton et al 2004). Interestingly, the blue dorsal pattern of *M. helenor* exhibits geographic variations, with populations displaying various widths of the central blue bands, and shades of blue within those bands defining sub-species (Blandin 2007). These geographic variations are parallel to the variation in the closely-related species *Morpho achilles* Linnaeus observed in sympatry throughout all the Amazonian basin: significant convergences in both the dorsal pattern (Llaurens et al 2020) and the blue iridescence (Ledamoisel et al 2026) have been documented between these two species, consistent with the expectations of evasive mimicry. Nevertheless, evidence stemming from predation pressure exerted on these butterflies from the relevant predator communities is still completely lacking.

Here, we exploit the early-morning-habit of insectivorous birds visiting light-traps used to attract nocturnal insects to perform predation experiments on dummy butterflies placed on the white sheet of these traps, allowing to characterise the community of wild birds actually attacking Lepidoptera and the effect of their traits (i.e., foraging behaviour and diet) on their attack decisions. By capturing the behaviours of wild communities of insect-eating birds, this experiment allows to address the following questions: 1) Do natural predators selectively avoid conspicuous colour patterns associated with evasiveness? If so, 2) are locally occurring evasive prey avoided more readily than exotic ones? To this aim, we compare predation rates exerted on prey that are (i) evasive i.e., prey harbouring *Morpho helenor* iridescent blue dorsal wing pattern (local and exotic form), (ii) unpalatable-control, i.e., prey harbouring the colour pattern of a local toxic aposematic prey (*Heliconius atthis* Doubleday), (iii) cryptic i.e., prey harbouring the colour pattern of the ventral cryptic wing coloration exhibited by the local *M. helenor* butterflies, and a (iv) palatable-control, i.e., prey harbouring the colour pattern of a local palatable cryptic prey (*Caligo bellerophon* Stichel). We mainly test three hypotheses: first that wild avian predators selectively avoid conspicuous colour patterns associated with high evasiveness, predicting lower predation rates on dummies harbouring the *M. helenor* dorsal blue pattern compared to the palatable control (*Caligo bellerophon*). Second, that this avoidance is driven by local frequency-dependent selection, therefore, dummies displaying the local *Morpho* phenotype will suffer fewer attacks than the exotic form. Lastly, we hypothesize that predation pressure depends on avian predatory traits, i.e. foraging or diet, so that defended prey will receive a higher proportion of attacks from generalist or opportunistic species than for specialists.

## MATERIALS AND METHODS

We performed a series of experiments in semi-wild conditions, using artificial butterflies pinned on insect light-traps that are visited by avian predators at dawn.

### Study site

Experiments were conducted in August 2024 and from July to August 2025 at four sites in northwest Ecuador (Table 1, supplementary material S1). Light traps at these reserves, which are active year-round (except in JQR), were used for the study. These traps are primarily installed for bird-watching activities. The light traps were distributed across primary and secondary forest, and forest patches embedded in human-disturbed areas, encompassing premontane and cloud forest ecosystems. One trap was sampled per day (experimental session), with 3-day revisit intervals in 2024 and 7-day intervals in 2025.

**Table 1.** Study sites, site codes, location coordinates, number of experimental sessions conducted at each site and geographical distance between sites.

| Site | Code | Location | N experiment.<br>session | Distance [km] |  |  |
| --- | --- | --- | --- | --- | --- | --- |
|  |  |  |  | MILR | JQR | CTR |
| Milpe Bird Sanctuary Reserve | MILR | 0°01'50"N 78°51'58"W | 17 |  |  |  |
| Reserva Jardín del Quinde | JQR | 0°02'08"N 78°52'19"W | 17 | 1 |  |  |
| Choco Toucan Reserve | CTR | 0°02'12"N 78°52'18"W | 18 | 1.2 | 0.2 |  |
| Reserva Amagusa-Mashpi | AMR | 0°09'35"N 78°51'11"W | 17 | aprox. 14 |  |  |

### Dummies

To test predator behaviour regarding prey harbouring different colour patterns, we built artificial dummies composed of wings of actual butterflies (supplementary material S2). Five different types of dummies were manufactured: (1) iridescent blue dorsal wing side of the local subspecies of *M. helenor* (*evasive-local*) (2) iridescent blue dorsal wing side of exotic subspecies of *M. helenor* (*evasive-exotic*), (3) brown ventral wing side of the local sub-species of *M. helenor* (*cryptic*) (4) dorsal wing pattern of *Caligo bellerophon* (*palatable-control*) and (5) dorsal wing patterns of *Heliconius atthis*, a locally-abundant butterfly harbouring high concentration of repellent cyanogenic glucosides (Pinheiro de Castro et al 2019) (*unpalatable-control*). The two subspecies of *Morpho helenor* (*evasive-local* and *evasive*-*exotic*) were used to quantify the effect of predator learning on the association between evasiveness and colour patterns. The local subspecies harbours an almost fully blue dorsal wing colour pattern, and is distributed along the west side slope of the Ecuadorian Andes, where the experiments were performed. The exotic subspecies presents an Amazonian distribution and harbours a dorsal transversal blue band on a black background, a wing pattern likely novel to birds participating in our experiments.

All butterflies were supplied by a local butterfly breeding farm (Quinta de Goulaine).

### Experimental set up

Experiments were conducted using pre-existing insect light traps (see supplementary materials <u>S2</u> for details and <u>S3</u>for a video) already installed at the reserves for bird-watching, where wild birds have become habituated to finding abundant insect prey (mostly Lepidoptera, Coleoptera and Hemiptera) each morning. No additional trapping or lighting infrastructure was installed for our research purposes. The light traps consist of a vertically hanging white sheet (dimensions: 2 by 2 m), illuminated by a 50-watt white LED bulb. Bird species attracted to the traps were primarily insectivores, although some frugivorous and granivorous species were also observed (see supplementary material S4). Two cameras (GoPro Hero 12) were used to capture a wide view of birds’ attacks. Individual bird species identity was always recorded during observations; however, individual birds could not be distinguished across attack events, and therefore repeated visits by the same individual could not always be identified. As a consequence, attack events were treated as observational units within sessions. In addition, two observers *in situ* (∼5 m from the light trap), equipped with binoculars, identified bird species and recorded the time and order of attacks when possible, allowing cross-validation with video recordings. When the experiment was finished, all dummies and cameras were removed from traps.

We simultaneously presented five different types of prey (dummies): evasive (1) *local* and (2) *exotic* (3) *cryptic*, (4) *palatable-control*, and (5) *unpalatable-control*. Dummies were pinned on the white sheet before bird activity began (around 5:15 am) interspersed among insects (hereafter referred to as alternative prey) attracted during the night to the light-trap. Dummy positions were randomized per experimental session. Observations generally started at 5:50 am and lasted one hour and 10 minutes (4200 s), corresponding to the peak period of bird activity. In six of the 69 experimental sessions, observations started a few minutes later because of adverse weather conditions, but the total observation duration remained the same across all sessions. We recorded the time and order of each attack event during experimental sessions. For dummy types that were not attacked during the observation period, attack latency was assigned a value of 4200 s (corresponding to the total duration of the experiment), and attack order was set to the number corresponding to the last attack of the session.

### Characterization of the local bird community

Bird species were identified using video recordings from the predation experiment. Given the scarcity of studies on Neotropical bird diets (Cadena-Ortíz et al 2025; Díaz et al 2024; Duca et al 2023), we defined only three feeding behaviour categories following Billerman et al (2026) (Fig. 1): (i) *flight feeding*, defined as aerial prey capture from a perch (e.g., Tyrannidae), or by following army ants to catch insects flushed by the swarms (e.g., Thamnophilidae); (ii) *terrestrial*, for birds foraging on or within the soil and leaf litter; and (iii) *treehunter*, for birds extracting food from bark (e.g., Furnariidae), branches or foliage (e.g., Thraupidae). Diet was assigned following the trophic niches proposed in Pigot et al (2020): frugivore, granivore, nectarivore, invertivore (consumers of invertebrates), carnivore. When two food types were assigned for a bird species in Billerman et al (2026), a combination of categories was used, starting with the most-consumed one.

**Fig 1.**
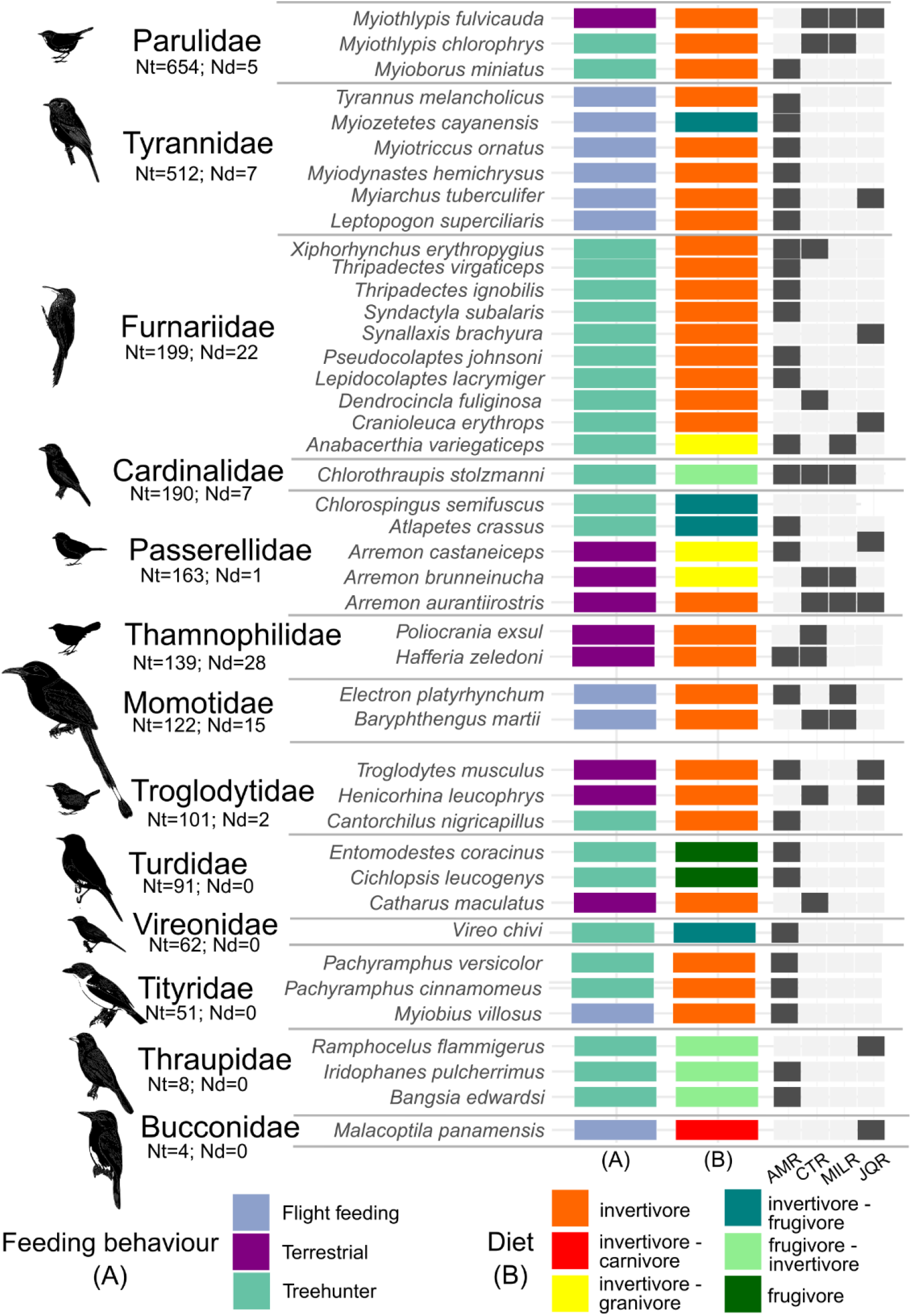
Bird species occurrences at each site (AMR, CTR, MILR and JQR) ranked by number of attack events. Black/grey tiles indicate presence/absence. Coloured tiles indicate feeding behaviour (A) or diet (B). Total attacks (Nt) and dummy attacks (Nd) are also shown.

### Statistical analyses

#### Attack latency and relative order of attack

To test whether evasive prey were attacked less frequently or less readily than toxic and cryptic prey, we compared predation responses among dummy categories using two metrics: (i) attack latency, defined as the time elapsed (in seconds) from the beginning of the experiment and the attack directed at a given dummy type, and (ii) relative attack order, calculated as the proportion of alternative prey items attacked before the dummy was attacked and relative to the total number of prey items (including actual live insects resting on the white sheet) attacked within a session. For unattacked dummy types, attack latency was set to 4200 s (total experiment duration), and relative attack order was set to 1. To avoid non-independence arising from repeated attacks on the same dummy type within a session, only the first attack recorded for each dummy type was retained for these analyses. This ensured that each prey type contributed a single observation per experimental session. Because individual birds could not be uniquely identified across attack events, predator identity could not be included in the statistical models. Analyses were therefore conducted at the level of prey type within experimental sessions. Differences among dummy types were tested using Kruskal–Wallis tests followed by Dunn’s post hoc pairwise comparisons with adjustment for multiple testing.

#### Attack probabilities depending on bird’s predatory traits

We calculated the log-likelihood of observing the events of attacks recorded on each dummy (*palatable-control, cryptic, evasive-local, evasive-exotic*, or *unpalatable-control prey*), compared to others within each category of birds’ predatory traits: (1) feeding behaviour (flight-feeding, terrestrial, tree-hunter) and (2) main diet (frugivore, invertivore), following the method implemented in previous studies (Mérot et al 2015; Willmott et al 2017; Páez et al 2021 and Linke et al 2022, 2025):

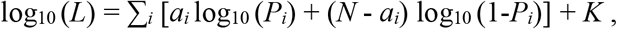

where *i* represents the dummy type; *N* is the total count of attack events on dummies; a*i* is the number of times a dummy of type *i* was attacked; *Pi* is the attack probability for each dummy type *i*; and *K* is a constant that disappears when comparing models. For each predatory category, we compared six scenarios to explore where attack rates of different dummy types could be equal or not (Table 2) and calculated the log-likelihood functions of those scenarios. Models were selected on the basis of their AICc: the model with the lowest AICc was selected as the best, while any model within 2 units of AICc was not rejected. We calculated Akaike weights following Burnham and Anderson (2002).

**Table 2.**
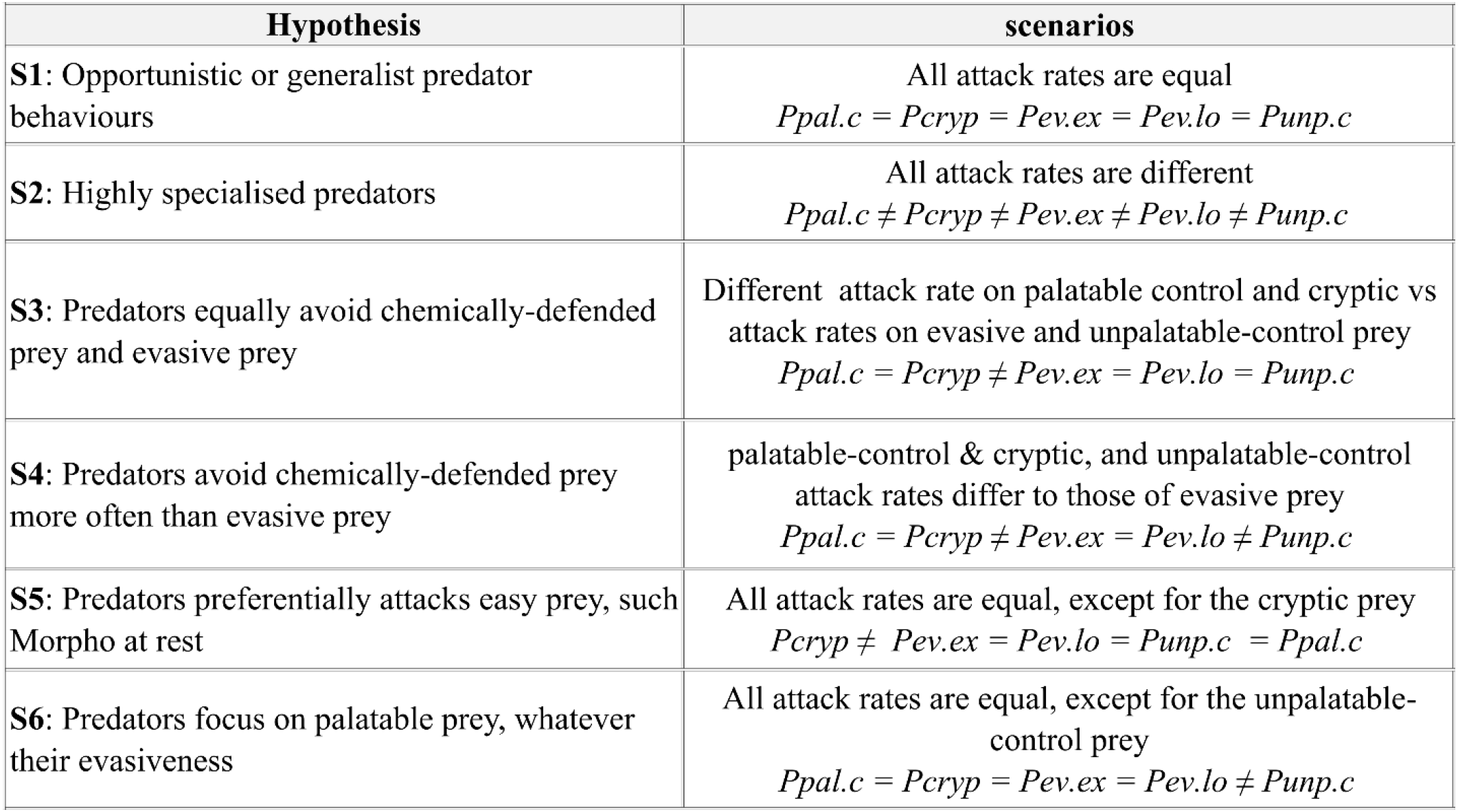
Scenarios investigated based on attack rates from the experiment considering feeding behaviour and diet of birds. *P* (probability) *pal*.*c:* palatable control; *cryp*: cryptic; *ev*.*ex*: evasive exotic; *ev*.*lo*: evasive local; *unp*.*c*: unpalatable control.

Statistical analyses (except log-likelihood tests, which were implemented in a spreadsheet) were done using R v.4.4.2 (R Core Team 2024). Graphics were generated using the R package ggplot2 v. 3.4.3 (Wickham 2016)

## RESULTS

### Local bird predator community

We observed attacks performed by 43 bird species across the 4 sampled sites, with varied feeding behaviour and diet (Fig. 1). Invertivorous species (invertivore, invertivore-granivore, invertivore-frugivore and invertivore-carnivore diet) were generally performing more attacks on our experimental sheets (37 out of 43), while frugivorous species (frugivore & frugivore-invertivore diet) (6 out of 43) were more scarcely observed.

### Predation on the different types of prey

Overall, both the time elapsed before striking the dummy (attack latency) and the rank of the stricken dummies (relative order) revealed the same trend (Fig. 2a,b). The *cryptic* prey was attacked first, and the *unpalatable-control* tended to be attacked much later or after a large number of alternative preys were consumed, as expected. The *palatable-control* also tended to be attacked very early during the experiments, but later than the *cryptic* prey. The two dummies displaying the dorsal blue pattern of *M. helenor* i.e., *evasive*, were attacked much later, at a time or a rank much closer to those of the *unpalatable-control*. The *evasive-exotic* pattern was attacked slightly earlier as compared to the *evasive-local*, (means secs: 3586, SD 1008 vs 3635, SD 979), although these differences were non-significant (*Post-hoc* Dunn’s test local vs exotic: z = 0.206, p = 0.836 [attack latency]; z = -0.310, p = 0.757 [relative order]) notably because of the limited number of attacks on our dummies (full results provided in supplementary material S5).

**Fig 2.**
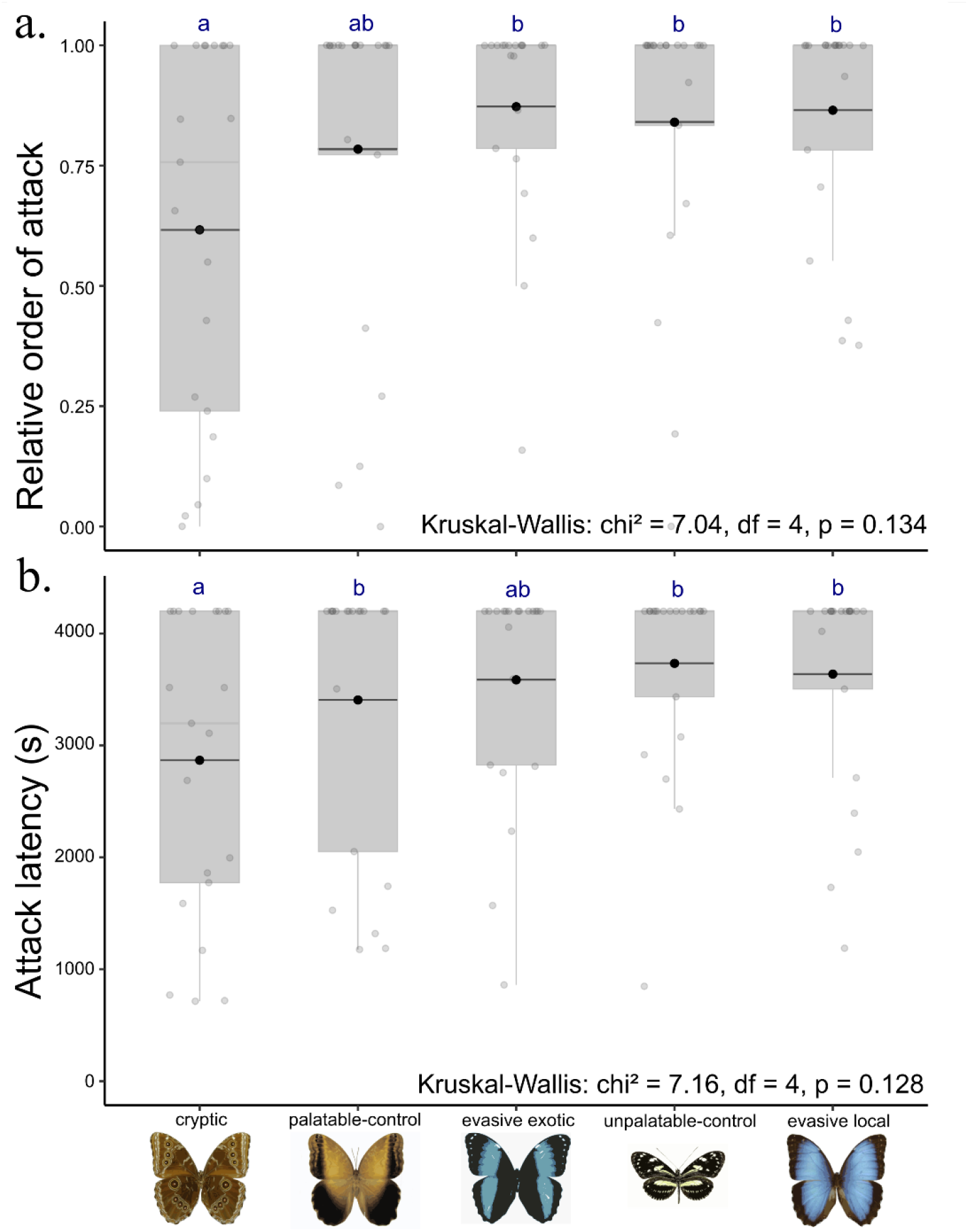
Comparison of relative order of attacks (a) and attack latency (b) on the different types of prey. Results from the Kruskal-Wallis and Post-hoc Dunn’s Test (pairwise comparison) are shown as well. Significant differences are indicated with letters in blue bold. Points at the upper extreme of the boxplot correspond mainly to no attacks.

### Attack behaviours of birds with different diet and foraging habits

Attack rates on dummy prey varied among birds with different feeding behaviours and diets (Fig. 3).

**Fig 3.**
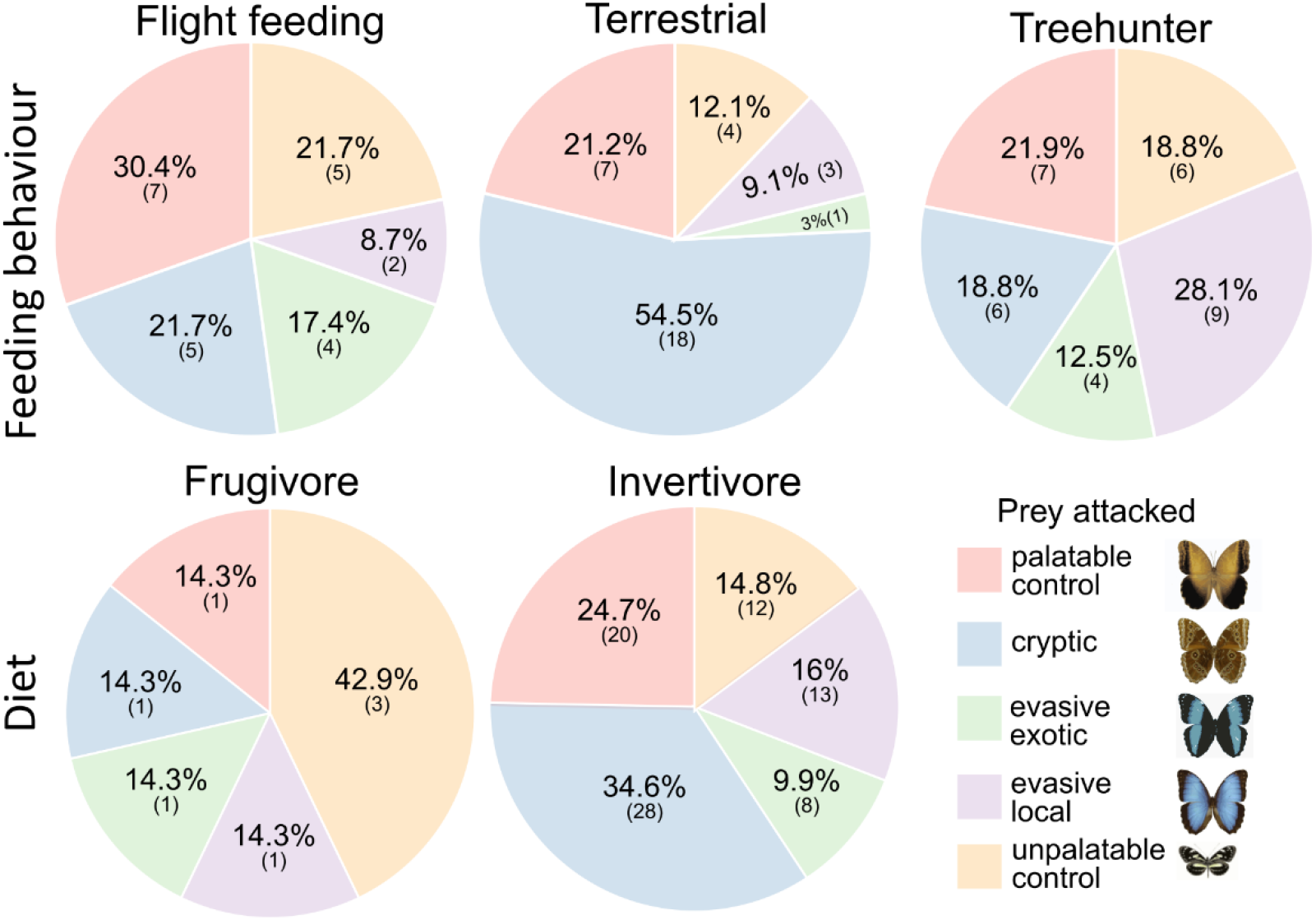
Proportion of observed bird attacks by prey type for each feeding behaviour and diet category.

These differences were assessed using likelihood-based models. The frugivore guild was excluded from the analysis due to the low number of observed attacks (N = 7). Our likelihood-based model test supported three main scenarios (Table 3): (i) no preference or avoidance for any type of prey i.e., opportunistic or generalist behaviour (S1) as the best scenario for flight-feeding and treehunter birds. (ii) Reduced attacks on conspicuous colour patterns (S3) as the best scenario for the invertivore birds, and second-best scenario (within 2 AICc units of the best model) for the flight feeding and terrestrial birds. (iii) cryptic prey as the most attacked among others (S5) as the best scenario for terrestrial, and second-best scenario for the invertivore birds. See supplementary material S6 for complete results.

**Table 3.**
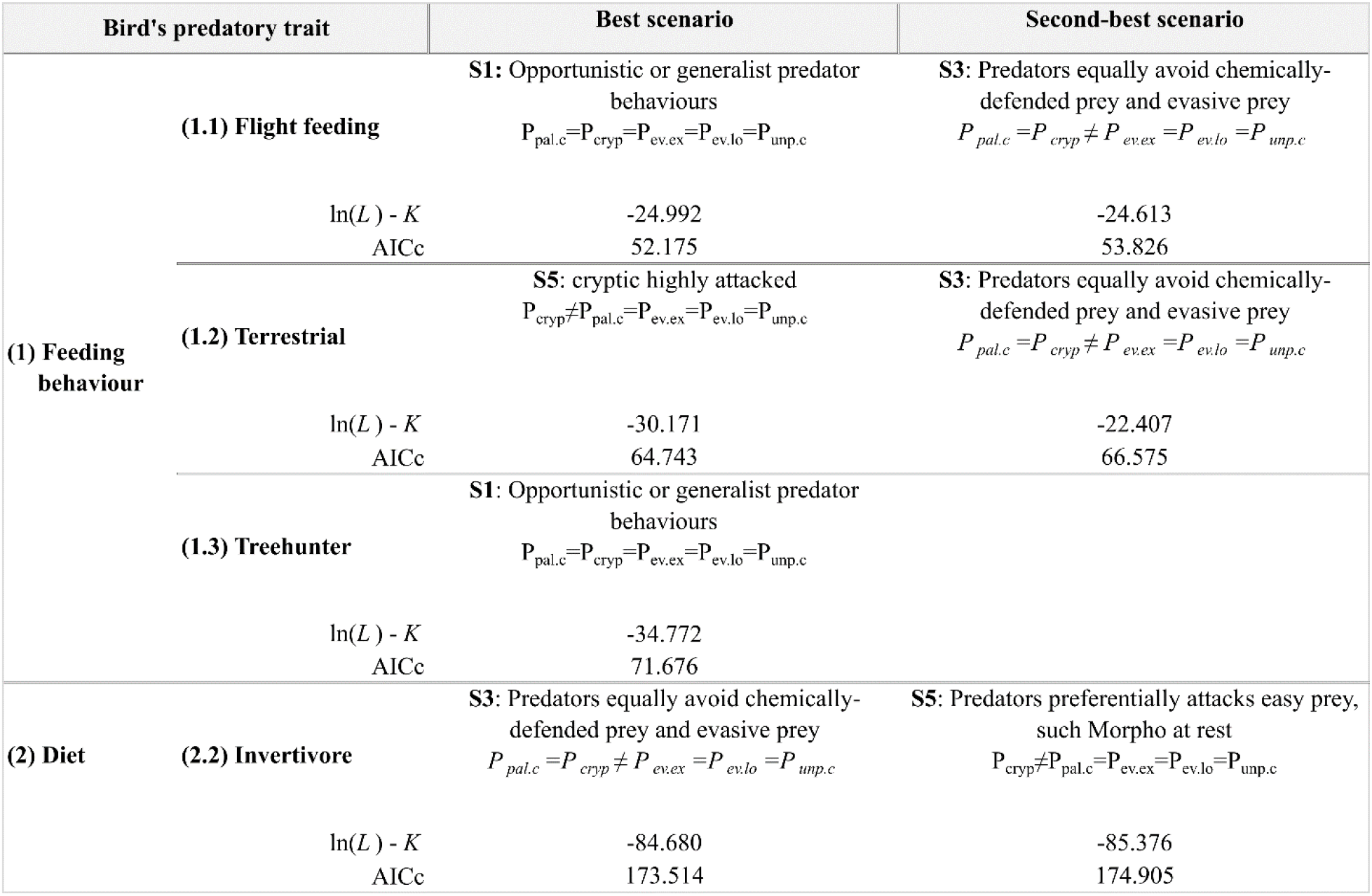
Best scenarios from observed attack rates among bird’s feeding behaviour and mainly diet based on the AICc value. Scenarios within a two-unit AICc interval with that of the best scenario are considered plausible as well.

## DISCUSSION

### A diverse avian predator community as a selective agent

To better understand the evolution of aposematism and mimicry in butterflies, it is crucial to study the behaviour of predators as they generate the main selective pressure acting on warning signals variation (Brower 1984; Mappes et al 2005). Despite extensive empirical approaches investigating selection acting on aposematic and mimetic signals using artificial or alive butterflies as prey and birds as predators (Benson 1972; Chai 1986; Gamberale-Stille and Guilford 2004; Pinheiro et al 2014; Wiklund and Järvi 1982), most studies have focused on a limited number of predator species. However, in nature, butterfly communities are exposed not to a single predator species, but to diverse avian assemblages exhibiting a wide range of foraging behaviours (Endler and Mappes 2004; Pinheiro and Cintra 2017; Willmott et al 2017). Greater predator diversity likely broadens the spectrum of visual capabilities and cognitive biases acting on prey survival chances (Endler and Mappes 2004). Consistent with this, our study shows that wild prey experience predation from a diverse avian community with differing foraging behaviours and diets, leading to differences in predation pressure among prey types.

### Conspicuousness in defended vs. palatable prey

Our observations align with extensive empirical research demonstrating that conspicuous signals often experience lower predation rates (Mappes et al 2005; Sillén-Tullberg 1985) and higher hesitation time (Schuler and Roper, 1992; Skelhorn et al 2016) compared to cryptic or dull patterns. In our experiment, camouflaged patterns presented on a white background were attacked more frequently and with shorter hesitation times than aposematic prey. We also noticed that the *evasive* wing patterns of *Morpho* butterflies were attacked at the same rate than the *unpalatable-control Heliconius* pattern, suggesting that prey evasiveness might provide similar defence against natural predators. This supports growing evidence for the evolution of aposematic signalling and mimicry in palatable but hard-to-catch prey (Pinheiro and Campos 2019; Le Roy et al 2019; Linke et al 2022, 2025; Páez et al 2026). Evasive signals may deter predators at least as effectively as chemical defences (Páez et al 2021): the cost of pursuing and missing a palatable evasive prey might be even larger than the cost of chasing and trying a slow but distasteful prey, leading to a faster avoidance of prey harbouring a colour pattern associated with high escape abilities. Conspicuousness in palatable species displaying high evasiveness can thus be under strong positive selection because it provides a significant protection against predator attacks.

But when considering the different predatory traits, this pattern stands only for specialists such as flight feeding, or invertivore birds. More opportunistic species such as treehunters or frugivore birds showed non-selective attack patterns. This underscores the importance of considering the full predator community, where heterogeneous foraging strategies impose contrasting selection pressures on prey defences and coloration.

### Evasive mimicry in *Morpho* butterflies

In this study, we hypothesized that local avian predators, being acquainted with the local *Morpho* pattern, would target the exotic pattern more frequently due to their lack of familiarity with the novel signal (Mallet and Barton 1989; Langham 2004; Chouteau et al 2016). However, our results showed no significant differences in avoidance rates or hesitation times, contradicting the expectation that novel signals incur higher predation. Neophobia i.e., avoidance towards novel stimuli (Greenberg and Mettke-Hofmann 2001) seems unlikely here given that attack rate on the *exotic* pattern was similar to the *unpalatable-contro*l prey. A plausible explanation could be broad predator generalisation (Páez et al 2021), potentially driven by the high energetic cost of pursuing evasive prey. In many classical mimetic systems, predators generalize based on highly-salient discriminatory traits such as intense coloration (Aronsson and Gamberale-Stille 2008; Rönkä et al 2018) which can overshadow other informative traits such as shape, patterns or size (Halpin et al 2013; Kazemi et al 2014). The iridescent blue coloration is not exclusive to *Morpho*. Other palatable yet evasive butterflies, such as members of the tribe Preponini and males of several *Doxocopa* Hübner (Apaturinae) species, are part of the “bright blue bands” ring described by (Pinheiro and Freitas 2014), a potential case of evasive mimicry. This suggests that the “blue iridescence” could be familiar as a visual cue within the natural predator community (although suggestive, Pinheiro and Freitas [2014] did not include *Morpho* as part of this or any other mimicry ring). Thus, in *Morpho helenor*, the iridescent blue, associated with erratic flap-gliding flight (Le Roy et al 2021) behaviour generating flash display (Murali 2018; Vieira-Silva et al 2024) likely acts as a salient cue that triggers avoidance at long distances (i.e. approx. 12 m (Pinheiro 1996; Pinheiro et al 2014; Pinheiro and Campos 2019) before predators can discriminate subtle differences between species. This is consistent with the theoretical framework proposed by Sherratt (2002), which predicts that predators may choose to avoid prey types that look remotely like highly unprofitable prey for fear of misclassification, even after learning is complete (Ruxton et al 2018). The protective effect of evasive mimicry could thus mainly rely on the highly salient trait i.e., blue iridescence associated with specific erratic flight behaviour, therefore relaxing selection and promoting accurate mimicry.

### Contrasted selection on ventral *vs*. dorsal wing patterns

Here, we observed that *Morpho* dummies displaying the cryptic ventral pattern received more attacks than those showing the conspicuous dorsal coloration by the flight-feeding predators. Terrestrial and treehunters accounted for a large proportion of attacks to the cryptic pattern, which may correspond to their behaviour regarding live *Morpho helenor* butterflies, which frequently rest on the ground with the wings folded when puddling or feeding, and might thus be more detected by non-specialist or opportunistic terrestrial and treehunter species. Many other butterflies exhibit conspicuous dorsal coloration with cryptic ventral surfaces (e.g., some Charaxinae species *Consul fabius* Cramer, *Archeoprepona demophon* Hübner, and many other species). The two sides are displayed in different ecological and behavioural contexts such as resting, basking, feeding or patrolling (Chai 1986; Pinheiro et al 2016; Pinheiro and Cintra, 2017; Pinheiro and Campos 2019). The contrasted evolution of dorsal *vs*. ventral colour pattern may result from antagonistic natural and sexual selective pressures (Puissant and Llaurens 2025) in different ecological contexts. Indeed, butterflies are likely more vulnerable to ambush predators when stationary (e.g., during feeding, puddling, or oviposition), whereas other predator guilds may dominate attacks during flight. In addition, Pinheiro et al. (2016) showed that experienced bird predators such as rufous-tailed Jacamars (the most specialized Neotropical butterfly predator (Chai 1986; Pinheiro 2011) sight rejects fast-flying butterflies like *Morpho*, while birds from other species, such as tyrant-flycatchers (Tyrannidae [Fitzpatrick 1980, 1981; Pinheiro 1996, 2003]), puffbirds (Bucconidae [Melo and Marini 1999]), motmots (Momotidae [Pinheiro et al 2008]), and anis (Cuculidae [Burger and Gochfeld 2001]), use alternative hunting tactics to locate and attack them when they stop flying and perch on a given substrate.

Finally, our results should be interpreted with caution. The low number of attacks limits statistical power, and the sampled bird community is likely biased towards early-active species attracted to light traps, thus representing only a subset of the local bird species assemblage. Nonetheless, this subset of the actual local bird communities still reveals substantial diversity in foraging and dietary traits, highlighting the importance of incorporating predator community structure into studies, rather than focusing on single “model” predators to characterize the selective landscape shaping the evolution of aposematism and mimicry in Neotropical systems.

## ETHICS STATEMENT

This study was performed under the permit issued by the Ministerio del Ambiente, Agua y Transición Ecológica MAATE-DBI-CM-2023-0298. This study was strictly observational and non-invasive in nature. No birds were captured, handled, restrained, marked, or collected. Behavioural observations were conducted using artificial models placed on pre-existing light traps independently operated by legally registered reserves as part of their routine ecotourism activities. No additional trapping or lighting infrastructure was installed for research purposes, and all procedures were designed to minimize disturbance to wildlife and surrounding habitats.

## ACKNOWLEDGEMENTS

This study was funded by the European Union (ERC-2022-COG - OUTOFTHEBLUE - 101088089). Views and opinions expressed are however those of the authors only and do not necessarily reflect those of the European Union or the European Research Council. Neither the European Union nor the granting authority can be held responsible for them. EPV’s postdoctoral fellowship was funded by Collège de France during this study. We thank the INABIO for providing permits for research in Ecuador, most recently under permit MAATE-DBI-CM-2023-0298. We thank Sergio Basantes and Doris Villalba from Reserva Mashpi-Amagusa, Jorge Naranjo from Reserva Ecológica Jardin del Quinde, Hans Heinz from Choco Toucan Reserve and Bird Lodge, Germania Cerezo, Liseth Simbaña, and Luis Yanes from Milpe Bird Sanctuary (Mindo Cloud Forest Foundation), Nikanor Mejia (Milpe), Mateo Roldán and Chiara Correa from Mashpi Lodge, Agustina Arcos and Alejandro Solano from Reserva-Finca Mashpi Shungo for allowing us to use their infrastructure for pilot and former experiments. We also thank Juan Freile, and Nicole Büttner for their advice during the early stages of the experimental design, and Mathieu de Goulaine for providing butterfly specimens. EPV thanks José Vazquez, José Pastrana, José Napa, Fabián Páez, Sebastián Mena, Alicia Arzani, Cristina Ríos and Martín Morocho for their assistance in the field.

## DATA ACCESSIBILITY STATEMENT

Raw data and R code that produce and support the findings of this study will be available in Zenodo repository upon publication.

## DECLARATION OF AI USE

We have not used AI-assisted technologies in creating this article.

