## Supplementary material S1 for "Do tropical birds avoid evasive prey? An experimental study in Ecuadorian avian communities"

**ELECTRONIC SUPPLEMENTARY MATERIALS**

**S1. Study site.** Map showing the four sites in northwest Ecuador where experiments were conducted in August 2024 and from July to August 2025. Milpe Bird Sanctuary Reserve (MILR [0°01'50"N 78°51'58"W; 1140 masl]), Jardín del Quinde Reserve (JQR [0°02'08"N 78°52'19"W; 1115 masl]), Toucan Chocó Reserve (CTR [0°02'12"N 78°52'18"W; 1116 masl]), and Amagusa-Mashpi Reserve (AMR [0°09'35"N 78°51'11"W; 1287 masl]). Sites are indicated with different colors. The points for MILR, JQR, and CTR overlap due to their close proximity in the general map but can be seen on the zoomed map. *Morpho helenor* colour patterns differ on either side of the Andes.

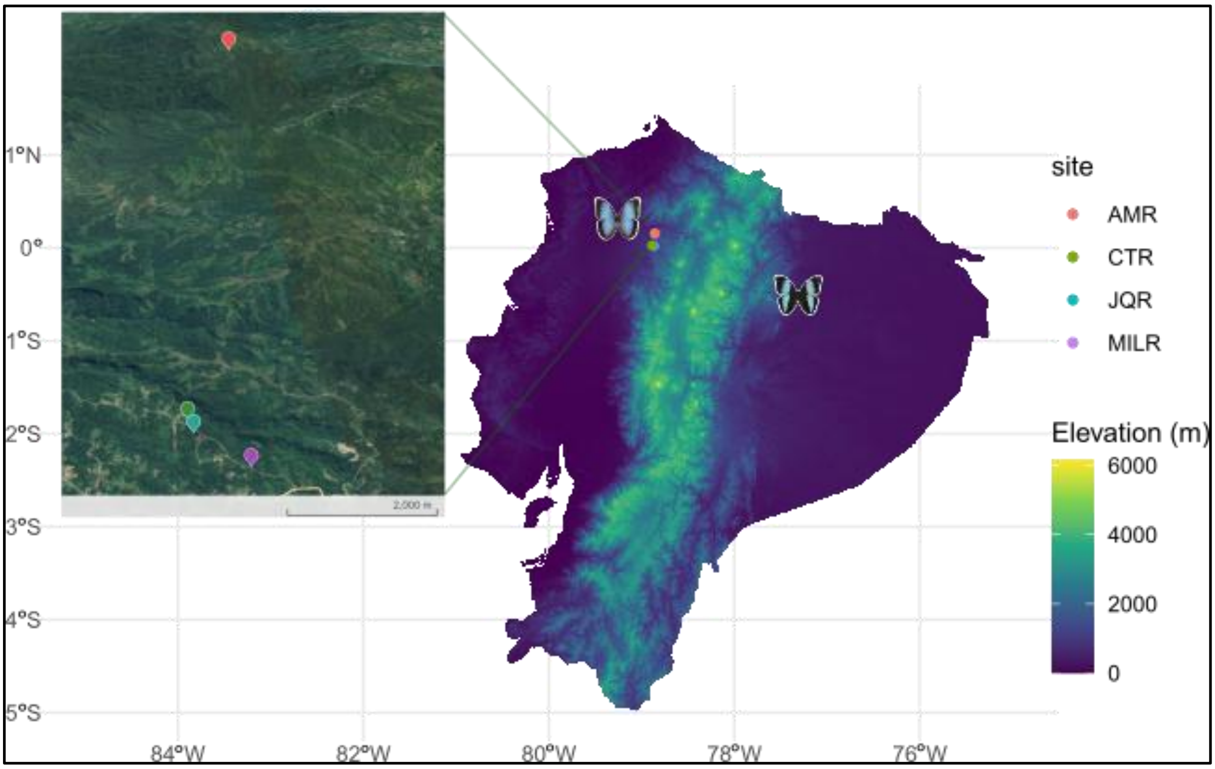
