## Supplementary material S2 for "Do tropical birds avoid evasive prey? An experimental study in Ecuadorian avian communities"

**S2. Experiment set up.** It consists of a light trap with naturally attracted insects (mostly moths) and artificial prey (dummies) manufactured with real wings of butterflies to test bird predation. Four dummies were placed at each corner of the sheet, with one in the center. Dummies positions were randomized at each experimental session. In the image (below) red arrows show the dummies: *Morpho* local (1), *Morpho* exotic (2), cryptic (3), palatable control - *Caligo* (4), unpalatable control - *Heliconius* toxic prey (5). One camera (GoPro Hero 12) was positioned perpendicularly and another at approximately 45 degrees in front of the light trap to capture a wide view of birds' attacks.

Two human observers, equipped with binoculars, were placed in a hidden spot, *i.e.*, a structure or shelter with viewing openings that conceal observers from birds, which allowed observers a close monitoring (approximately 5 m from the light trap) without disturbing the birds.

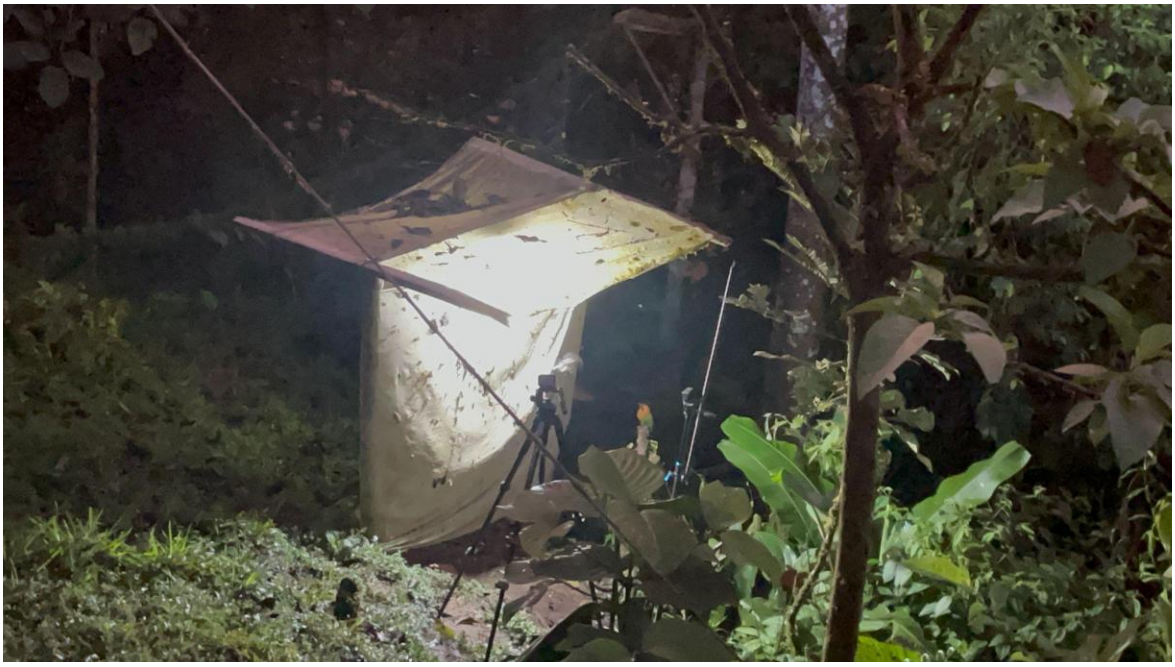

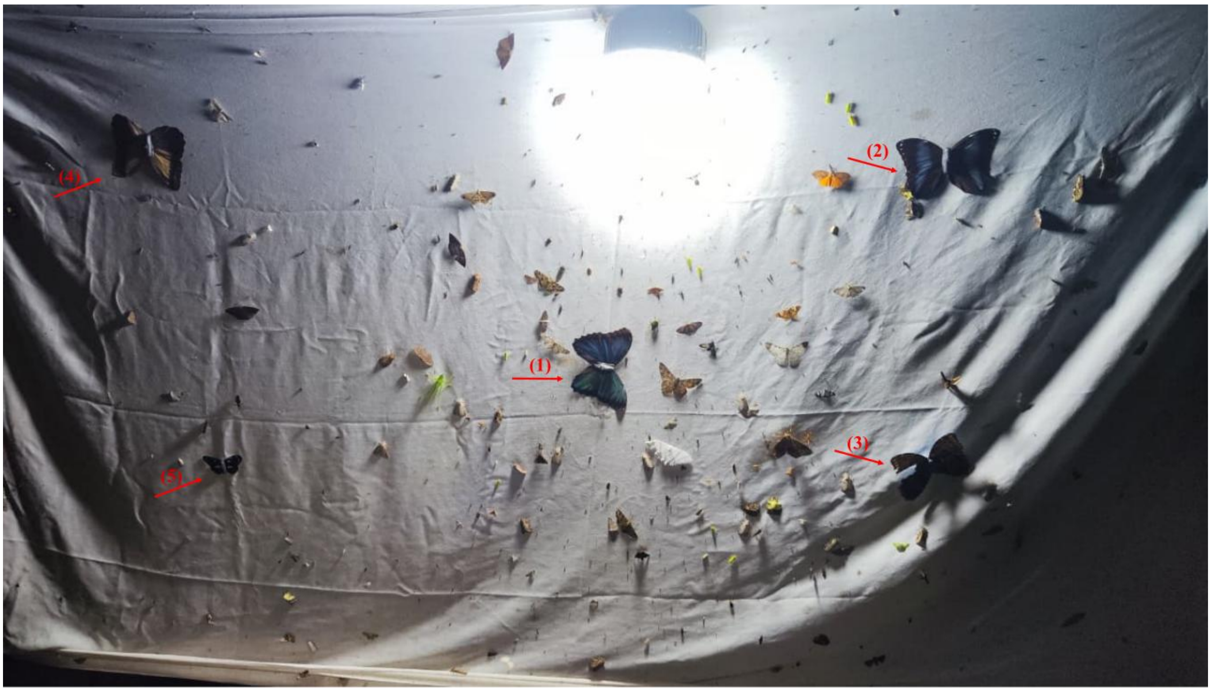
(1)
(2)
(3)
(4)
(5)
