## Supplementary material S4 for "Do tropical birds avoid evasive prey? An experimental study in Ecuadorian avian communities"

**S4. List and distribution of bird species attracted to light-traps during our study.** Birds that consume two types of food, are presented as a combination of these categories, starting with the one that is consumed the most. Reserva Amagusa-Mashpi (AMR), Choco Toucan Reserve (CTR), Milpe Bird Sanctuary Reserve (MILR) and Reserva Jardín del Quinde (JQR).

| Species | Feeding behaviour | Diet | Site | | | |
|---|---|---|---|---|---|---|
| | | | AMR | CTR | MILR | CTR |
| *Anabacerthia variegaticeps* | treehunter | invertivore | * | | * | |
| *Arremon aurantiirostris* | terrestrial | invertivore-granivore | | * | * | * |
| *Arremon brunneinucha* | terrestrial | invertivore-granivore | | * | * | |
| *Arremon castaneiceps* | terrestrial | invertivore-granivore | * | | | |
| *Atlapetes crassus* | treehunter | invertivore-frugivore | * | | | |
| *Bangsia edwardsi* | treehunter | frugivore-invertivore | * | | | |
| *Baryphengus martii* | flight feeding | invertivore | | * | * | |
| *Cantorchilus nigricapillus* | treehunter | invertivore | * | | | |
| *Catharus maculatus* | terrestrial | invertivore | | * | | |
| *Chlorospingus semifuscus* | treehunter | invertivore-frugivore | | | | * |
| *Chlorothraupis stolzmanni* | treehunter | frugivore-invertivore | * | * | * | |
| *Cichlopsis leucogenys* | treehunter | frugivore | * | | | |
| *Cranioleuca erythrops* | treehunter | invertivore | | | | * |
| *Dendrocincia fuliginosa* | treehunter | invertivore | | * | | |
| *Electron platyrhynchum* | flight feeding | invertivore | * | | * | |
| *Entomodestes coracinus* | treehunter | frugivore | * | | | |
| *Hafferia zeledoni* | terrestrial | invertivore | * | * | | |
| *Henicorhina leucophrys* | terrestrial | invertivore | | * | | * |
| *Iridophanes pulcherrimus* | treehunter | frugivore-invertivore | * | | | |
| *Lepidocolaptes lacrymiger* | treehunter | invertivore | * | | | |
| *Leptopogon superciliaris* | flight feeding | invertivore | * | | | |
| *Malacoptila panamensis* | flight feeding | invertivore-carnivore | | | | * |
| *Myarchus tuberculifer* | flight feeding | invertivore | * | | | * |
| *Myiobius villosus* | flight feeding | invertivore | * | | | |
| *Myioborus miniatus* | treehunter | invertivore | * | | | |
| *Myiodynastes hemichrysus* | flight feeding | invertivore | * | | | |
| *Myiothlypis chlorophrys* | treehunter | invertivore | | * | * | |
| *Myiothlypis fulvicauda* | terrestrial | invertivore | | * | * | * |
| *Myiotriccus ornatus* | flight feeding | invertivore | * | | | |
| *Myiozetetes cayanensis* | flight feeding | invertivore-frugivore | * | | | |
| *Pachyramphus cinnamomeus* | treehunter | invertivore | * | | | |
| *Pachyramphus versicolor* | treehunter | invertivore | * | | | |
| *Poliocrania exsul* | terrestrial | invertivore | | * | * | |
| *Pseudocolaptes johnsoni* | treehunter | invertivore | * | | | |
| *Ramphocelus flammigerus* | treehunter | frugivore-invertivore | | | | * |
| *Synallaxus brachyura* | treehunter | invertivore | | | | * |
| *Syndactyla subalaris* | treehunter | invertivore | * | | | |
| *Thripadectes ignobilis* | treehunter | invertivore | * | | | |
| *Thripadectes virgaticeps* | treehunter | invertivore | * | | | |
| *Troglodytes musculus* | terrestrial | invertivore | * | | | * |
| *Tyrannus melancholicus* | flight feeding | invertivore | * | | | |
| *Vireo chivi* | treehunter | invertivore-frugivore | * | | | |
| *Xiphorhynchus erythropygius* | treehunter | invertivore | * | * | | |
