## Supplementary material S5 for "Do tropical birds avoid evasive prey? An experimental study in Ecuadorian avian communities"

**S5. Results from the Kruskal-Wallis and Post-hoc Dunn's Test (pairwise comparison).** Attack latency metric was calculated by the time in seconds elapsed from the beginning of the experiment to the first attack per session to the dummy. Relative order metric was calculated by the number of other preys attacked before the strike on the dummy divided by the total number of attacks per session. Significant results are in bold font.

| | **Relative order of attack** | | **Attack latency** | |
|---|---|---|---|---|
| **Type of prey** | mean | sd | mean | sd |
| cryptic | 0.617 | 0.395 | 2348 | 1559 |
| palatable control | 0.784 | 0.360 | 3095 | 1610 |
| evasive exotic | 0.873 | 0.223 | 3196 | 1418 |
| unpalatable control | 0.865 | 0.229 | 3354 | 1338 |
| evasive local | 0.840 | 0.296 | 3393 | 1264 |
| **Kruskal-Wallis test** | chi$^2$ = 7.039, df = 4, p = 0.134 | | chi$^2$= 7.062, df = 4, p = | |
| **Post-hoc Dunn's Test** | Z | *p* | Z | *p* |
| palat.control - cryptic | 1.907 | 0.056 | 1.719 | 0.857 |
| palat.control- evasive exotic | -0.045 | 0.964 | -0.084 | 0.933 |
| cryptic - evasive exotic | -1.953 | **0.051** | -1.803 | 0.072 |
| palat.control - evasive local | -0.355 | 0.722 | -0.579 | 0.563 |
| cryptic - evasive local | -2.263 | **0.024** | -2.298 | **0.022** |
| evasive exotic - evasive local | -0.310 | 0.757 | -0.494 | 0.621 |
| palat.control - unpalat.control | -0.256 | 0.798 | -0.537 | 0.592 |
| cryptic - unpalat.control | -2.163 | **0.031** | -2.256 | **0.024** |
| evasive exotic - unpalat.control | -0.210 | 0.833 | -0.452 | 0.651 |
| evasive local - unpalat.control | -0.099 | 0.921 | 0.042 | 0.966 |
