## Supplementary material S6 for "Do tropical birds avoid evasive prey? An experimental study in Ecuadorian avian communities"

| | | | Hypothesis | S1: Opportunistic or generalist predator behaviours | S2: Highly specialised predators | S3: Predators equally avoid chemically-defended prey and evasive prey | S4: Predators avoid chemically-defended prey more often than evasive prey | S5: Predators preferentially attacks easy prey, such Morpho at rest | S6: Predators focus on palatable prey, whatever their evasiveness |
|---|---|---|---|---|---|---|---|---|---|
| | | | Scenario | Ppal.c = Pcryp = Pev.ex = Pev.lo = Punp.c | Ppal.c ≠ Pcryp ≠ Pev.ex ≠ Pev.lo ≠ Punp.c | Ppal.c = Pcryp ≠ Pev.ex = Pev.lo = Punp.c | Ppal.c = Pcryp ≠ Pev.ex = Pev.lo ≠ Punp.c | Pcryp ≠ Pev.ex = Pev.lo = Punp.c = Ppal.c | Ppal.c = Pcryp = Pev.ex = Pev.lo ≠ Punp.c |
| **Flight Feeding** | | | | | | | | | |
| | attacked | presented | attack rates | | | | | | |
| palatable-control | 7 | 23 | *Ppal.c* | 0.2 | 0.304 | 0.261 | 0.261 | 0.196 | 0.196 |
| cryptic | 5 | 23 | *Pcryp* | 0.2 | 0.217 | 0.261 | 0.261 | 0.217 | 0.196 |
| evasive-exotic | 4 | 23 | *Pev.ex* | 0.2 | 0.174 | 0.159 | 0.130 | 0.196 | 0.196 |
| evasive-local | 2 | 23 | *Pev.lo* | 0.2 | 0.087 | 0.159 | 0.130 | 0.196 | 0.196 |
| unpalatable-control | 5 | 23 | *Punp.c* | 0.2 | 0.217 | 0.159 | 0.217 | 0.196 | 0.217 |
| | 23 | 115 | | | | | | | |
| number of parameters (attack rates) | | | | 1 | 5 | 2 | 3 | 2 | 2 |
| ln(L) - K | | | | -24.992 | -24.164 | -24.613 | -24.432 | -24.980 | -24.980 |
| AICc | | | | **52.175** | 61.858 | **53.826** | 56.127 | 54.561 | 54.561 |
| **Terrestrial** | | | | | | | | | |
| | attacked | presented | attack rates | | | | | | |
| palatable-control | 7 | 33 | *Ppal.c* | 0.2 | 0.212 | 0.379 | 0.379 | 0.114 | 0.220 |
| cryptic | 18 | 33 | *Pcryp* | 0.2 | 0.545 | 0.379 | 0.379 | 0.545 | 0.220 |
| evasive-exotic | 1 | 33 | *Pev.ex* | 0.2 | 0.030 | 0.081 | 0.061 | 0.114 | 0.220 |
| evasive-local | 3 | 33 | *Pev.lo* | 0.2 | 0.091 | 0.081 | 0.061 | 0.114 | 0.220 |
| unpalatable-control | 4 | 33 | *Punp.c* | 0.2 | 0.121 | 0.081 | 0.121 | 0.114 | 0.121 |
| | 33 | 165 | | | | | | | |
| number of parameters (attack rates) | | | | 1 | 5 | 2 | 3 | 2 | 2 |
| ln(L) - K | | | | -35.858 | -28.886 | -31.088 | -30.864 | -30.171 | -35.477 |
| AICc | | | | 73.845 | 69.994 | **66.575** | 68.555 | **64.743** | **75.354** |
| **Treehunter** | | | | | | | | | |
| | attacked | presented | attack rates | | | | | | |
| palatable-control | 7 | 32 | *Ppal.c* | 0.2 | 0.219 | 0.203 | 0.203 | 0.203 | 0.203 |
| cryptic | 6 | 32 | *Pcryp* | 0.2 | 0.188 | 0.203 | 0.203 | 0.188 | 0.203 |
| evasive-exotic | 4 | 32 | *Pev.ex* | 0.2 | 0.125 | 0.198 | 0.203 | 0.203 | 0.203 |
| evasive-local | 9 | 32 | *Pev.lo* | 0.2 | 0.281 | 0.198 | 0.203 | 0.203 | 0.203 |
| unpalatable-control | 6 | 32 | *Punp.c* | 0.2 | 0.188 | 0.198 | 0.188 | 0.203 | 0.188 |
| | 32 | 160 | | | | | | | |
| number of parameters (attack rates) | | | | 1 | 5 | 2 | 3 | 2 | 2 |
| ln(L) - K | | | | -34.772 | -34.207 | -34.770 | -34.763 | -34.763 | -34.763 |
| AICc | | | | **71.676** | 80.721 | 73.954 | 76.383 | 73.940 | 73.940 |
| **Frugivore** | | | | | | | | | |
| | attacked | presented | attack rates | | | | | | |
| palatable-control | 1 | 7 | *Ppal.c* | 0.2 | 0.143 | 0.143 | 0.143 | 0.214 | 0.143 |
| cryptic | 1 | 7 | *Pcryp* | 0.2 | 0.143 | 0.143 | 0.143 | 0.143 | 0.143 |
| evasive-exotic | 1 | 7 | *Pev.ex* | 0.2 | 0.143 | 0.238 | 0.143 | 0.214 | 0.143 |
| evasive-local | 1 | 7 | *Pev.lo* | 0.2 | 0.143 | 0.238 | 0.143 | 0.214 | 0.143 |
| unpalatable-control | 3 | 7 | *Punp.c* | 0.2 | 0.429 | 0.238 | 0.429 | 0.214 | 0.429 |
| | 7 | 35 | | | | | | | |
| number of parameters (attack rates) | | | | 1 | 5 | 2 | 3 | 2 | 2 |
| ln(L) - K | | | | -7.063 | -7.063 | -7.499 | -7.063 | -7.565 | -7.063 |
| AICc | | | | **18.013** | 84.126 | 21.999 | 28.126 | 22.130 | 21.126 |
| **Invertivore** | | | | | | | | | |
| | attacked | presented | attack rates | | | | | | |
| palatable-control | 1 | 7 | *Ppal.c* | 0.2 | 0.247 | 0.296 | 0.296 | 0.164 | 0.213 |
| cryptic | 1 | 7 | *Pcryp* | 0.2 | 0.346 | 0.296 | 0.130 | 0.346 | 0.213 |
| evasive-exotic | 1 | 7 | *Pev.ex* | 0.2 | 0.099 | 0.136 | 0.130 | 0.164 | 0.213 |
| evasive-local | 1 | 7 | *Pev.lo* | 0.2 | 0.160 | 0.136 | 0.130 | 0.164 | 0.213 |
| unpalatable-control | 3 | 7 | *Punp.c* | 0.2 | 0.148 | 0.136 | 0.148 | 0.164 | 0.148 |
| | 7 | 35 | | | | | | | |
| number of parameters (attack rates) | | | | 1 | 5 | 2 | 3 | 2 | 2 |
| ln(L) - K | | | | -88.015 | -83.934 | -84.680 | -84.646 | -84.376 | -87.625 |
| AICc | | | | 178.081 | 178.557 | **173.514** | 175.604 | 174.905 | 179.403 |
